# Mechanotransduction of strain regulates an invasive phenotype in newly transformed epithelial cells

**DOI:** 10.1101/770487

**Authors:** Sophie Chagnon-Lessard, Hubert Jean-Ruel, Michel Godin, Andrew E. Pelling

## Abstract

Our organs and tissues are in constant motion, exposing epithelial cells to mechanical stretch. However, how these external forces impact cellular morphology, organization and dynamics in healthy and diseased tissues is still being elucidated. In several studies, we and others have shown how mechanical stresses and strains in the epithelium can modulate the dynamics and invasiveness of transformed cells. Carcinoma, the most common type of cancer, develops in the sheets of cells forming the epithelium and lining our organs and cavities. It usually begins with the transformation of a single cell via the activation of oncogenes such as Ras. Here, we show in a model system how mechanical stretch in epithelial sheets results in a more invasive phenotype in transformed cells.

Cyclic strain prevents the formation of strong circumferential belts of actin in Ras^V12^ cells and greatly promoting the formation of Ras^V12^ protrusions, typical of a more invasive phenotype. We also show that Ras^V12^ and wild type MDCK cells possess distinct sensitivity to strain. External forces remodel their actin cytoskeletons and adhesion complexes differently, resulting in a more invasive system dynamic. Our work demonstrates that the Rho-ROCK mechanotransduction pathway is involved in regulating a mechanically-induced switch to a more invasive phenotype. In a mechanically dynamic microenvironment, transformed cells exhibit drastically different cellular dynamics and movements when compared to static conditions. They grow larger invasive protrusions, potentially making them harder to be eliminated from healthy tissues. The insights gained in this study reveal the complex dynamics at play in healthy and transformed epithelial cells which are found in a mechanically active microenvironment.

## 1 Introduction

Although they are one of the simplest tissues present in complex multicellular organisms, epithelial mono-and bi-layers play a crucial role in first-line protection of organs and cavities they envelop [1,2]. Epithelial cells form tight physical barriers constituting an efficient defense against pathogens and preventing the passage of other cells and macromolecules[3]. The monolayers maintain their integrity and barrier function despite continuous cell division and death (normal epithelial cell turnover). Apical extrusion is the mechanism by which a dying cell is eliminated, but this cellular process is also part of normal cell competition [4–6]. Certain cells exhibiting abnormal activities, such as some oncogenic expression, are also apically extruded, although in a death-independent manner [7,8]. The majority of human cancers result from the transformation of a single cell following alteration of its genome [9]. Mutations converting a gene belonging to the Ras family into an active oncogene is found in 20% of human tumors [10]. Ras proteins influence numerous signaling pathways, such as Rho [11,12], leading to changes in actin cytoskeleton configuration as well as, among others, cell shape, contractility, adhesion, motility, and division. Previous studies showed that H-Ras^V12^-transformed cells can be recognized and eliminated via apical extrusion by the concerted action of the surrounding wild type (WT) cells [8,13,14]. This has been referred to as epithelial defense against cancer or transformed cells. Diversion of the extrusion direction from apical to basal is observed in certain oncogenic cells, and is a potential mechanism to initiate metastasis [7,15–17]. Although the majority of Ras^V12^ cells are apically eliminated, some initiate basal invasions by remaining in the monolayer and growing protrusions. In certain cancers, the prominence of protrusions has been linked to invasiveness[18]. Importantly, these extrusion and protrusion behaviors of Ras^V12^-transformed cells are only observed when surrounded by WT cells, showing that heterotypic interactions play a crucial role.

Cells are constantly sensing and responding to mechanical forces and physical properties of the surrounding extracellular matrix (ECM) [19]. Through tissue matrix deformations and cytoskeleton-adhesion interfaces, most cells in vivo are stretched or compressed to a certain degree [20], for instance via muscle contraction, breathing, pumping of the heart, fluid pressure, or digestion. In this context, understanding the mechanical contribution to tumor initiation and development is required for a holistic comprehension of cancer biology [21,22]. Many studies investigate how the ECM stiffness impacts tumor progression [23], but mechanically dynamic microenvironments are considered less often. Two recent studies illustrate the importance of integrating mechanical factors to our understanding of cancer cell behavior. They showed that tugging forces on human fibrosarcoma cells grown on a 3D collagen/fibronectin scaffold lead to invadopodia maturation and increased cell invasion [24,25].

For the recognition and elimination of transformed cells, epithelial defense relies on many cytoskeletal proteins and cell-generated contractile forces in both WT [14] and transformed cells [26,27]. Given that external forces can remodel the cytoskeleton architecture, we hypothesize that external forces have a profound impact on the behavior of Ras^V12^ cells within the epithelial monolayer. To test this hypothesis, we fabricated polydimethylsiloxane (PDMS) microfluidic stretchers to mimic the complex non-uniform strain field occurring in vivo. We used this lab-on-a-chip strategy to mechanically stretch a cellular model system composed of Madin-Darby Canine Kidney (MDCK) H-Ras^V12^-WT co-cultures. MDCK cells are epithelial cells from the kidney tubules, which are small structures playing an important role in filtering blood. They are unavoidably subjected to small mechanical deformations, similarly to other compliant structures that operate under fluid pressure such as blood vessels [28]. We found that the application of external strains triggers Ras^V12^ cellular basal invasion, in part by activating the Rho-ROCK signaling pathway. Their apical extrusion is significantly reduced while protrusion formation is greatly promoted. We demonstrated that Ras^V12^ and WT cells have mismatched mechanoresponses with distinct cytoskeletal reorganization, shifting the system dynamic.

## 2 Results

### 2.1 External mechanical forces promote Ras^V12^ invasiveness

To generate a biologically relevant strain field, we employed an all-PDMS pneumatic-based microfluidic stretcher which we reported previously [29]. Cells were cultured on a thin suspended membrane encased in a micro-chamber. This membrane was cyclically stretched via the deformation of the cell chamber walls, upon the activation of adjacent vacuum chambers (see Movie S1 and Fig. 1B,C). The MDCK H-Ras^V12^-WT (1:75) cellular system was co-cultured on the type-I collagen-treated membrane (relaxed state), and incubated in a tetracycline-free environment until a monolayer was fully formed (10 h). Cyclic stretching (1 Hz, 3 to 9 % strain amplitude) and Ras^V12^ expression (induced by addition of tetracycline) were then simultaneously initiated. The systems were imaged after 24 hours of stretching. A set of automated image analysis programs was developed to precisely characterize the strain-induced changes of key cellular components (in particular e-cadherin and actin) throughout the co-culture. Fig. 1A-C present the working principle of the device and a portion of the image analysis.

**Fig 1.**
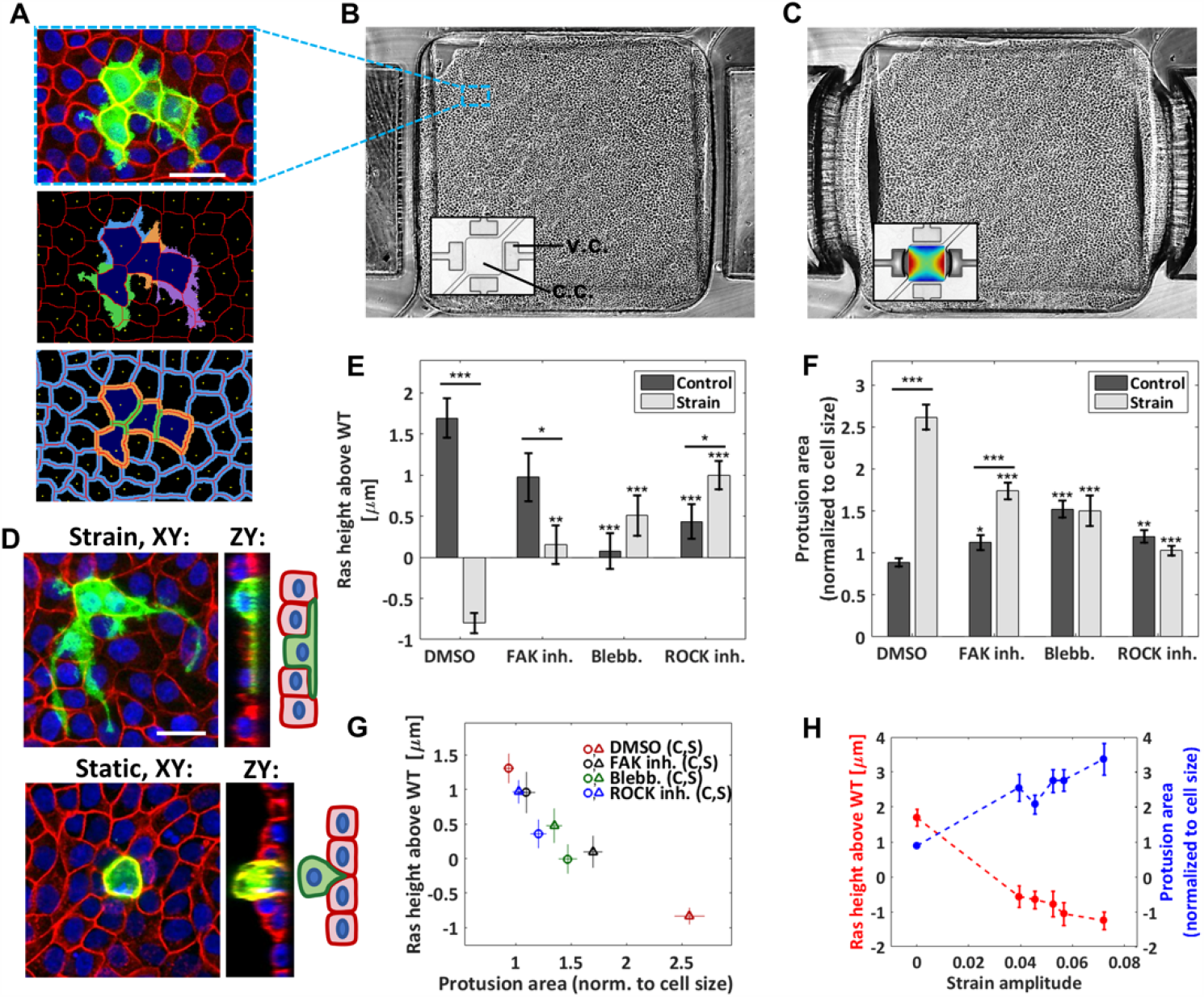
Cyclic stretching increases protrusion formation and hinders apical extrusion. **(A)** Top: confocal image of the MDCK co-culture (Ras^V12^ and WT cells, static condition) with GFP-Ras^V12^ cells in green, actin in red, and nuclei in blue. Scale bar is 35 μm. Middle and bottom: image analysis output showing the nucleus centers (yellow), the cellular contours (red), the Ras^V12^ cell bodies (dark blue), associated protrusions (middle) and the junctional regions (bottom: Ras^V12^-Ras^V12^ in green, Ras^V12^-WT in orange, and WT-WT in blue). **(B, C)** Monolayer grown in a PDMS pneumatic-based microfluidic stretcher in its relaxed (B) and stretched (C) states (phase contrast mode). The insets (bright field mode) show the empty device with the strain map superimposed in the stretched case (dark blue: 3%, dark red: 9%). The vacuum and cell chambers are identified as v.c. and c.c., respectively. The cell chamber width is 1.6 mm. **(D)** Confocal images of GFP-Ras^V12^ transformed cells in WT monolayers 24 h after tetracycline addition followed by F-actin (red) and nuclei (blue) staining. Scale bar is 25 μm. Top: representative example of Ras^V12^ cell forming large protrusions obtained in a stretched experiment. Bottom: representative example of extruding Ras^V12^ cell obtained in a static control. **(E)** Average height difference between the Ras^V12^ and WT cells for the static controls and stretched experiments. **(F)** Average protrusion area relative to the cell body (i.e. excluding the protrusions) for the static controls and the stretched experiments. The drug-free case (DMSO) and three drugs were tested: FAK inhibitor, blebbistatin, and ROCK inhibitor. Data are mean ± s.e.m. *** P<0.0001, **P<0.005, *P<0.05. Stars above the individual bars relate to the corresponding DMSO condition. Stars above the horizontal lines refer to the significance between the control and strain data. From left to right in (E, F): n = 234, 188, 78, 139, 268, 233, 237 and 334 cells each from 3 or 4 independent experiments. **(G)** Correlation between the average Ras^V12^ cell height and the average protrusion area for the 8 conditions presented in (E, F). **(H)** Average Ras^V12^ cell height (red) and average protrusion area (blue) as a function of the strain amplitude, for the DMSO condition. The zero-strain points correspond to the average of the control data, and the non-zero strain points were obtained by binning the data of the stretched experiments into five bins each containing an equal (37 or 38) number of cells.

We first examined the Ras^V12^ cell behavior within the WT monolayer when no strain was applied within the microfluidic device. We observed the formation of small basal protrusions and found that a significant fraction of the Ras ^V12^ cells were extruding, as expected [13]. An example of an extruding cell is shown in the lower image of Fig. 1D. Fig. 1E shows that their average height was above that of the surrounding WT cells (Supplementary Material Fig. S1). To study the impact of external mechanical forces on the system, we then examined the Ras^V12^ cell behavior under cyclic stretching and we observed a striking effect. Firstly, the average height of the Ras^V12^ cells dropped below that of the WT cells (Fig. 1E), suggesting a preponderance of basal – rather than apical – extrusion. Secondly, the average size of basal protrusions increased significantly (Fig. 1F), indicating increased aggressiveness in our model system. A representative example of a Ras^V12^ cell growing long protrusions under cyclic stretching is shown in the upper image of Fig. 1D. The dependence on strain amplitude of the average protrusion size and Ras^V12^ cell height indicates that the degree of invasiveness is modulated by the force amplitude (Fig. 1H). As the resulting variation is small in our system, data were simply grouped as either strain or static elsewhere. Overall, our results demonstrate that biologically relevant strains (3 to 9 %) trigger an invasive Ras^V12^ cell phenotype and impede the monolayer’s ability to eliminate transformed cells.

Pharmacological treatments were used to interfere with the mechanical stability of cells. Blebbistatin and ROCK inhibitor Y-27632, which respectively inhibit myosin II activity and actomyosin contractility, both decreased Ras^V12^ extrusion under static conditions (Fig. 1E). This agrees with previous observations which demonstrated the importance of ROCK in the apical extrusion process[13]. Under cyclic stretching, both blebbistatin and ROCK inhibitor significantly reduced the effects of strain on Ras^V12^ cell behavior (Fig. 1E,F). This indicates that myosin II activity and actomyosin contractility are involved in the increased invasiveness induced by strain. We found that inhibiting focal adhesion kinase (FAK) with PF-573228 also decreased the effect of strain, albeit more moderately (despite having the strongest impact on cell shape (Fig. S2). FAK stabilizes the actin cytoskeleton via the ROCK pathway and regulates central proteins in focal adhesions[30], thus playing a key role in mechanosensing and transduction[31–33]. FAK inhibition is being investigated as a new therapeutic cancer target[22,34] and has been shown to bypass some extrusion defects[16]. Interestingly, it is only when stretching is applied that we observe a reduction of Ras^V12^ invasiveness upon FAK inhibitor addition. These results show that the structural integrity of the cytoskeleton strongly affects the process by which substrate strains change the system dynamics and trigger Ras^V12^ invasive behavior.

In consideration of all conditions tested (DMSO, blebbistatin, ROCK and FAK inhibitions, each combined to both static and cyclically stretched substrates), we found a strong correlation between the average protrusion size of the Ras^V12^ cells and their average height relative to the surrounding WT cells (Fig. 1G). The decrease in apical extrusion is consistent with the facilitated protrusion growth of Ras^V12^ cells under strain. From a mechanistic point of view, it is more difficult for the WT cells to “squeeze out” a transformed cell which has increased adhesive contact area with the substrate, as hinted by Gudipaty *et al*.[1]. Conversely, reduced extrusion may facilitate protrusion progression.

### 2.2 Cortical actin belts around Ras^V12^ cells diminish under strain

The process by which epithelial monolayers expel dying cells to maintain their integrity is now believed to begin with the formation of a cortical contractile F-actin belt in the dying cell itself[4]. The cortical actomyosin cytoskeleton of a cell produces contractile forces that are coupled to neighboring cells via e-cadherin adhesions[2,27,35]. Specific patterns of intercellular tension are controlled in part by the cortical F-actin network. Dysregulation of this pattern of contractility at the cell-cell junctions has been shown to drive oncogenic extrusion[26,27]. An increase in cortical contractile F-actin within the Ras^V12^ cells was also found to be actively implicated in their extrusion by the surrounding WT cells[26,27]. Since these previous results indicate that cortical actin is critical in determining the fate of Ras^V12^ cells, we investigated the effects of strain on the remodeling of cortical actin throughout the co-culture. We observed, in agreement with previous studies[13,26], the presence of a strong cortical actin belt in Ras^V12^ cells under static conditions (Fig. 2A,B). In contrast, upon cyclic stretching (in the drug-free case), the difference in cortical actin intensities between Ras^V12^ and WT cells vanishes. This suggests that external strains at least partially alter the apical extrusion mechanism by preserving uniformity among the actomyosin contractile patterns in Ras^V12^ and WT cells. Upon the addition of FAK inhibitor, blebbistatin, and ROCK inhibitor, the strain-induced alteration of the actin ratio is largely suppressed. That is, the formation of a strong actin belt is not eliminated (Fig. 2B and Fig. S3). This shows that the abolition of the strong actin belt via external strains is at least partially regulated by the Rho-ROCK pathway. It also further supports the importance of the actin belt in the extrusion process, as highlighted by the strong correlation between the junctional actin intensity ratio Ras^V12^-Ras^V12^/WT-WT and the extruding Ras^V12^ height (Fig. 2C). Importantly, these results show that strain remodels the cytoskeleton differently in Ras^V12^ and WT cells, specifically at their junction, thus affecting their interaction.

**Fig 2.**
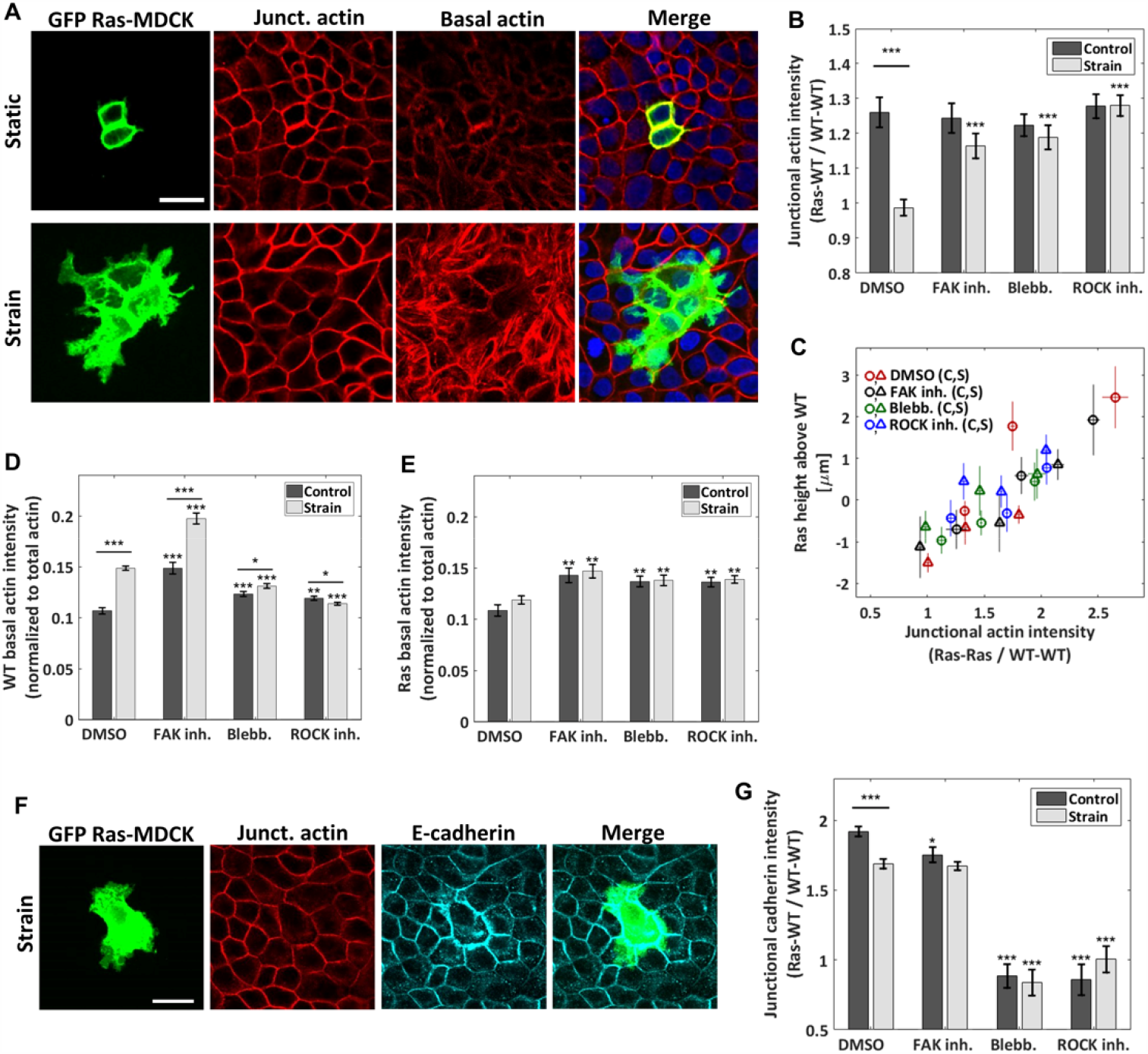
Cyclic stretching promotes actin and e-cadherin reorganization differently in WT and in Ras^V12^-transformed cells. **(A)** Confocal images of the Ras^V12^-WT culture for a static control (top) and a stretched experiment (bottom). GFP-Ras^V12^ cells are in green. Junctional and basal actin images (red) were obtained as described in the *Materials and Methods* and *Supplementary Material* S2 sections. Scale bar is 25 μm. **(B)** Ratio of the junctional actin intensity, Ras^V12^-WT interface over WT-WT interface. **(C)** Correlation between Ras^V12^ extrusion height and the junctional actin intensity ratio Ras^V12^-Ras^V12^ over WT-WT. The data from each of the 8 conditions (DMSO and three drugs, static (c) and strain (s)) were separated in three bins each of equal number of cells. **(D, E)** Average basal actin intensity of the WT (D) and Ras^V12^ (E) cells. Both are normalized to the actin intensity summed over all stacks and averaged over the full image. **(F)** Representative confocal images of the Ras^V12^-WT culture under strain showing a GFP-Ras^V12^ cell (green), junctional actin (red) and e-cadherin (magenta). Scale bar is 25 μm. **(G)** Ratio of the e-cadherin junctional intensity, Ras^V12^-WT interface over WT-WT interface. For (B, D, E, G), three drugs were tested in addition to the drug-free case (DMSO): FAK inhibitor, blebbistatin, and ROCK inhibitor. Data are mean ± s.e.m. *** P<0.0001, **P<0.005, *P<0.05. Stars above the individual bars relate to the corresponding DMSO control or DMSO strain condition. Stars above the horizontal lines refer to the significance between the control and strain data. From left to right, n = 43, 48, 29, 44, 56, 56, 53, and 57 images, each from 3 to 4 independent experiments. The average ratios of each image were compiled and they were used to determine the mean and the s.e.m. of the data points reported; the total number of Ras^V12^ cells contained in each set of images is given in Fig. 1E,F.

### 2.3 Depletion of the e-cadherin adhesive belt in Ras^V12^ cells under strain

Previous studies have shown that the loss of e-cadherin is an important step in the development of cancer[36] and that the loss of cell-cell adhesions is related to more invasive phenotypes[37,38]. For instance, activating RhoA and ROCK was found to promote cancer cell invasion via the disruption of e-cadherin adherent junctions[39]. We investigated the effects of strain on junctional e-cadherin signal intensity (Fig. 2F,G, Fig. S4 and S5). We observed that strains induce changes in e-cadherin differently in WT than in Ras^V12^ cells. In the static control, the accumulation of e-cadherin at Ras^V12^-WT (or Ras^V12^-Ras^V12^, Fig. S3) interfaces is greater than that at the WT-WT interfaces. This e-cadherin intensity mismatch diminishes upon cyclic stretching. This relative depletion of the e-cadherin adhesive belt in Ras^V12^ cells is coherent with a model in which increased RhoA-ROCK activity, in our case via mechanical strains, promotes a more invasive phenotype.

### 2.4 Strain promotes basal SF formation in WT cells but not in Ras^V12^ cells

To further investigate the mechanical landscape in which Ras^V12^ cells evolve, we analyzed the basal actin network of the monolayer. Almost no basal actin stress fibers (SFs) were visible in the static DMSO control (Fig. 2A). In contrast, upon stretching, they were largely promoted in the WT cells as shown in Fig. 2A,D. Indeed, the activation of the Rho-ROCK pathway (and the subsequent increase in myosin phosphorylation) following the application of substrate strain on cells promotes the formation of the actomyosin cytoskeleton[40]. Sahai *et al*.[41] showed that the generation of contractile forces via the Rho-ROCK pathway disrupts cell-cell junctions. From mechanical considerations, a well-developed actomyosin network exerting high tension tends to reduce cell-cell contact and adhesion strength[35,42]. This suggests that the WT-WT cell adhesion integrity is altered by the increased actomyosin network following cyclic stretching. Since the fate of Ras^V12^ cells depends on their interaction with the surrounding WT cells, we propose that this remodeling of the monolayer facilitates the progression of Ras^V12^ protrusions, since they grow preferentially at the WT-WT interfaces (see details below).

For both the drug-free condition and the three pharmacological treatments, no significant difference in Ras^V12^ basal actin was observed between the static and stretched conditions (Fig. 2A,E). This significant discrepancy with WT cells further demonstrates the mismatch in strain mechanoresponsiveness between Ras^V12^ and WT cells, and gives a hint that the former appear less responsive.

### 2.5 Strain-induced cellular reorientation is different in Ras^V12^ and WT cells

To further demonstrate the strain sensitivity mismatch between Ras^V12^ and WT cells we analyzed the cellular reorientation, a common signature of mechanoresponse upon cyclic-stretching. Although it is generally studied in elongated cells such as fibroblasts, it has also been observed in epithelial cells, including in MDCK cells[43,44]. Interestingly, we found that while WT cells preferentially reorient perpendicular to the stretching direction, Ras^V12^ cells exhibit a nearly random distribution (Fig. 3A). These results further support the interpretation that WT cells are more mechanoresponsive to cyclic stretching than Ras^V12^ cells (in the drug-free case). This reorientation mismatch was smaller (or non-existent) upon addition of pharmacological treatments. Fig. 3B shows a positive correlation between the reorientation mismatch (Ras^V12^ vs WT) and the degree of Ras^V12^ cell invasiveness for the four pharmacological conditions investigated (DMSO, FAK inhibitor, blebbistatin, and ROCK inhibitor). A greater reorientation mismatch coincides with larger basal protrusions and a lower average Ras^V12^ height. While we do not suggest causality between these parameters, it is consistent with our general results showing that Ras^V12^ and WT cells have a different sensitivity to strain. Additionally, we found a lower level of basal vinculin intensity in Ras^V12^ cells than in WT cells under cyclic stretching, suggesting differences in the connections between the substrate and F-actin networks [45] (Fig. S6).

**Fig 3.**
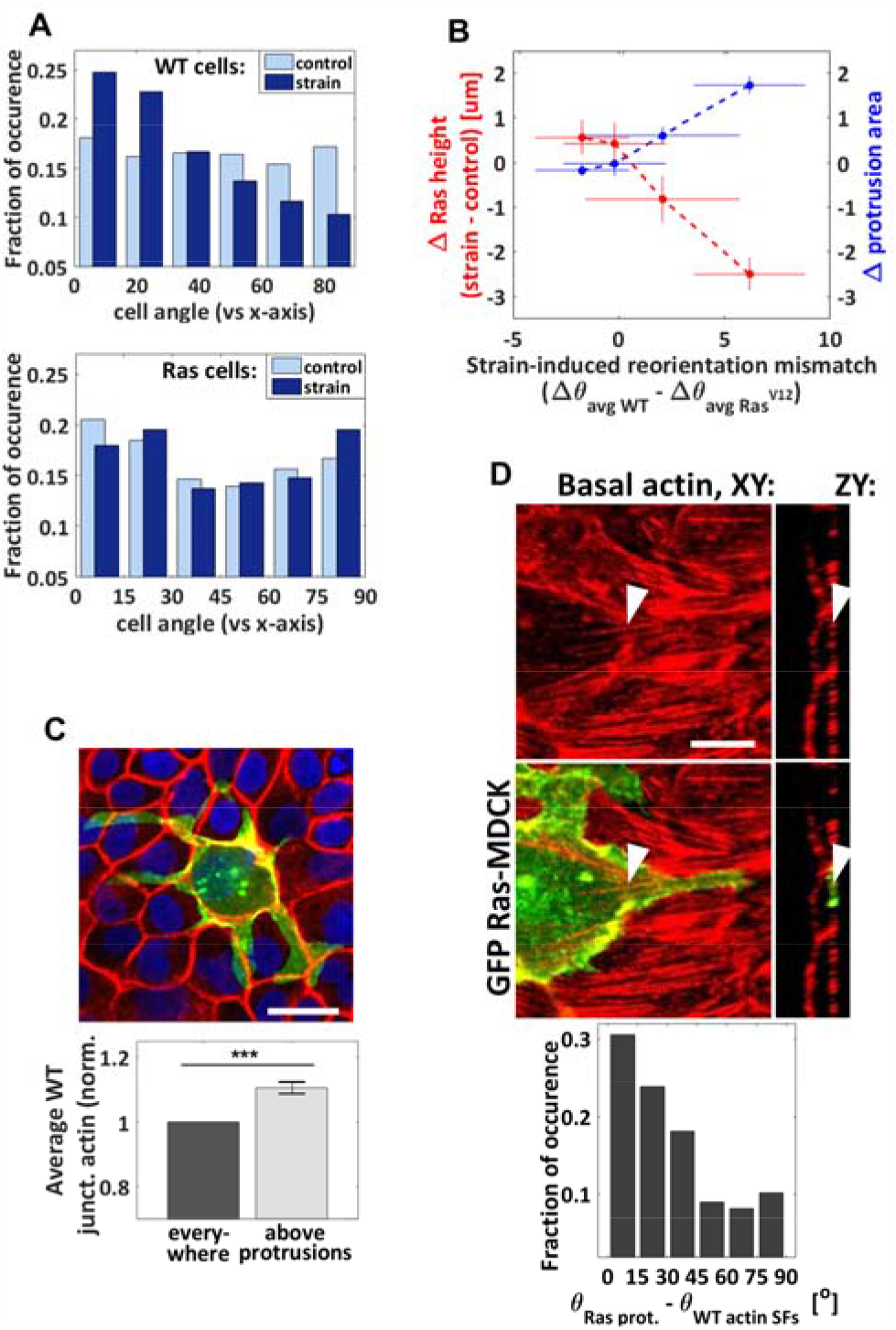
Orientation analysis. **(A)** Cellular reorientation response to cyclic stretching. Normalized incidence histogram of the angle difference between the cell body direction and the stretching axis for the WT (top) and Ras^V12^ (bottom) cells. n_WT control_ = 5261 cells, n_WT strain_ = 5076 cells, n_Ras control_ = 287 cells, and n_Ras strain_ = 189 cells from 3 independent strain and 3 independent control experiments. **(B)** Correlation between the reorientation mismatch and the strain-induced changes in average Ras^V12^ cell height and protrusion area. Each point corresponds to one pharmacological condition (from left to right: ROCK inhibitor, blebbistatin, FAK inhibitor, and DMSO). The reorientation mismatch is the difference between the strain-induced reorientation of the WT cells (Δθ *avg WT*) and that of the Ras^V12^cells (Δθ _*avg Ras*_), where Δθ _*avg*_ = θ _*avg strain*_ − θ_*avg control*_. When the mismatch is zero, the degree of cell reorientation under strain is the same for Ras^V12^ and WT cells. When it is higher than zero, the reorientation response is greater in WT cells. **(C)** Top: rather extreme example of Ras^V12^ cell with protrusions growing preferentially at the WT-WT junctions. Scale bar is 11 μm. Bottom: the average WT junctional actin intensity is greater above the Ras^V12^ protrusions than it is in average elsewhere. Control and strain DMSO conditions combined together, n = 91 images (each image contained 1 to 10 Ras^V12^ cells from 1 to 4 clusters, and hundreds of WT cells) from 3 independent control and 3 independent strain experiments. Data are mean ± s.e.m., *** P<0.0001. **(D)** Top: representative confocal images of basal actin (red) and GFP-Ras^V12^ (green) showing a protrusion growing parallel to neighboring actin SFs. The arrows point at the same SF in two different views. Scale bar is 6 μm. Bottom: normalized incidence histogram of the angle difference between the protrusions and the neighboring WT actin filaments (under strain only). This analysis could not be performed for the static condition due to the scarcity of basal actin filaments in the absence of stretching. n = 190 protrusion segments from 3 independent experiments.

### 2.6 The surrounding WT cells guide the Ras^V12^ protrusion growth direction

The ability of transformed cells to grow protrusions is known to facilitate and guide their invasion and progression [18,46,47]. We observed that the application of strain promotes protrusion formation in our model system, although it is as yet unclear what influences their growth direction. We analyzed their orientation with respect to the strain direction and found a nearly random distribution (not shown). Interestingly, we observed that protrusions grow preferentially parallel to the strain-induced basal SF network of the neighboring WT cells (Fig. 3D). As the driving force for protrusion growth is associated with the polymerization of localized actin filaments[46], it is possible that less resistance is encountered along that direction. Protrusion formation is thus guided at least partially by their interaction (either mechanical or chemical) with the surrounding WT cells, whose organization is modified by strain. We also found that Ras^V12^ cell protrusions grow preferentially along WT-WT interfaces (Fig. 3C). The progression of protrusion that seems facilitated under cyclic stretching (Fig. 1F) is thus consistent with the strain-induced promotion of the actomyosin contractile apparatus, demonstrated previously to weaken intercellular junctions[40]. These findings demonstrate that the structure of the WT monolayer modulates Ras^V12^ protrusion progression under cyclic stretching.

They suggest that the tissues comprising the microenvironement of transformed cells, and importantly the way they are shaped by physiological forces, are critical in guiding basal invasion.

## 3 Discussion

Cyclic stretching is known to induce actin cytoskeleton remodeling and increase actomyosin contractility via the activation of the Rho-ROCK signaling pathway, downstream phosphorylation of the myosin light chain kinase (MLCK), and myosin II activation [40]. It is also recognized that reorganization of the cytoskeleton is involved in the process of cancer cell invasion and metastasis [1,22,48]. A direct link has been established between the activation of the Rho signaling pathways (in particular targeting actin remodeling) and the ability of a tumor cell to initiate invasion [47,49]. RhoC and localized RhoA have been shown to play critical roles in protrusion growth[18,49]. Here we demonstrated that pharmacological treatments inhibiting ROCK or myosin II activity hinder the effects of strain on the system by partially restoring apical extrusion and inhibiting protrusion formation. This indicates that cyclic stretching drives the system toward Ras^V12^ invasiveness through the activation of cytoskeleton-modifying proteins, at least in part by activating the Rho pathway (see Fig. 4). Interestingly, while ROCK inhibition and ROCK activation (via strain) suppress apical extrusion when applied independently, their combined effects partially restore it. Since both of these conditions are known to interfere with ROCK in opposite ways, it suggests that a precise cytoskeleton mechanical configuration is required to enable apical extrusion.

**Fig 4.**
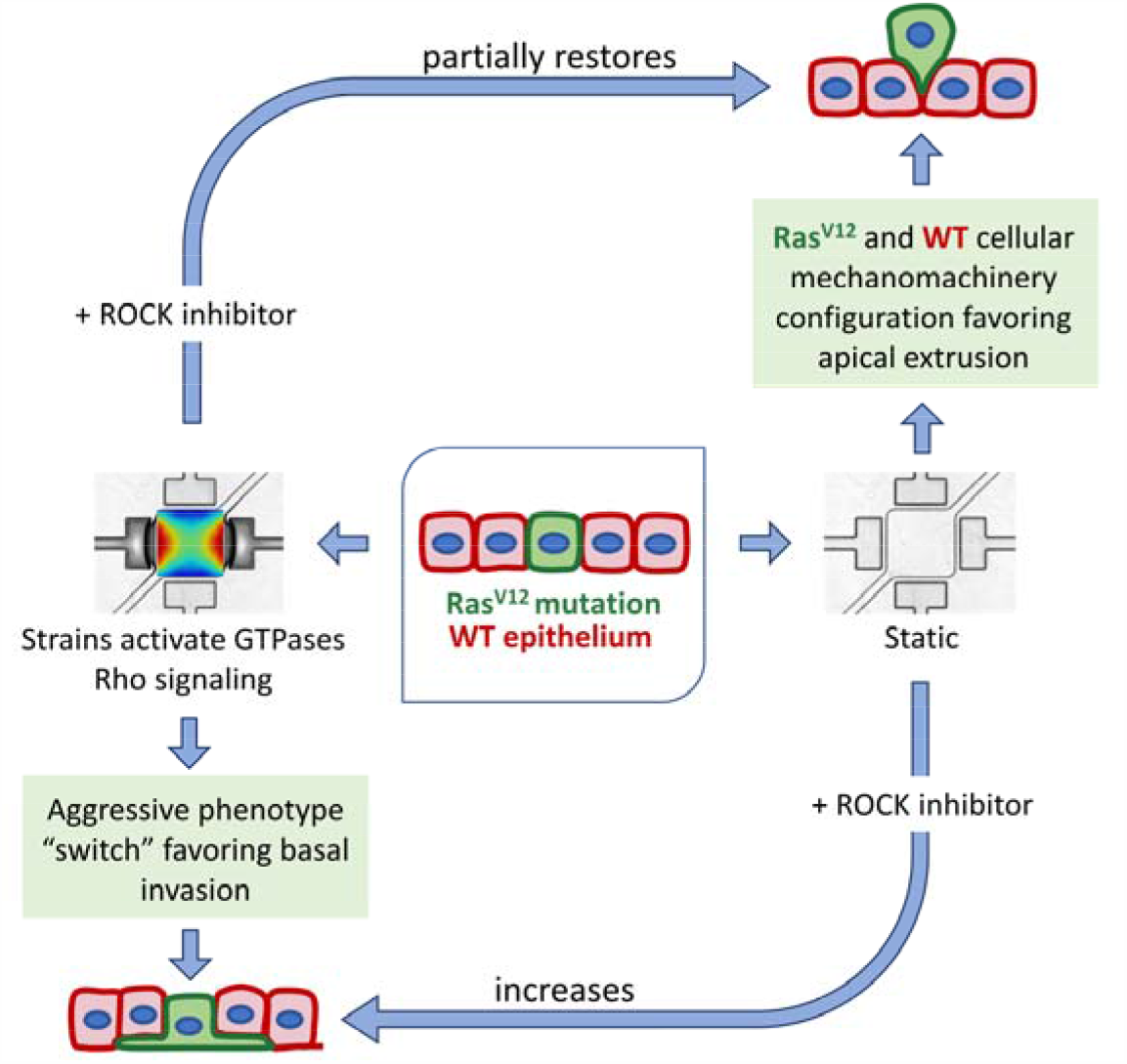
Scheme of the mechanically-induced switch to aggressive phenotype, showing the involvement of the Rho-ROCK mechanotransduction pathway. Under static condition, the specific Ras^V12^ and WT mechanomachinery configuration favors apical extrusion processes. A perturbation to this system, via mechanical stretching, abolishes the formation of the actin belt around Ras^V12^ cells and potentiate Ras^V12^ protrusion growth. The addition of ROCK inhibitor in a mechanically dynamic microenvironment partially restore the cortical actin belt around Ras^V12^ cells and the apical extrusion process.

The process by which transformed epithelial cells escape their primary site is not fully understood [7]. Apical extrusion is the typical mechanism by which epithelial cell turnover takes place. In contrast, oncogenic mutations can alter this process, resulting in basal extrusion, and potentially enabling invasion and escape towards the tissue they envelop [7,15–17]. For instance, K-Ras cells (not studied here) were shown to be predominantly basally extruded via degradation of S1P, preventing the formation of the strengthened contractile belt in the surrounding WT cells [15]. S1P is also known to be required for the H-Ras apical extrusion [50]. Its binding to S1PR2 receptors in WT cells leads to filamin accumulation (a force generator) via the Rho-ROCK pathway. Here we found that although apical extrusion is preponderant in H-Ras cells under static conditions, cyclic stretching drives cell extrusion basally. In light of this, it would be relevant to study the S1P levels in the system to determine if they are involved in the mechanism by which strain promotes basal invasion.

Furthermore, all the results described in this work were obtained using an impenetrable solid PDMS membrane as the substrate, which is different from in vivo conditions. Although the impact of external forces on Ras^V12^ cell invasiveness is unambiguous in our model system, it would be enlightening to use porous membranes in future studies to fully investigate the basal extrusion process. More generally, further work is needed since the type of cell, microenvironement, and mechanical stimulus greatly affect the cellular response. Increased invasion of human fibrosarcoma cells was observed in a collagen/fibronectin scaffold when subjected to tugging forces, in line with our results [24,25]. Interestingly, daily stretching exercises in mice was found to reduce in vivo tumor volume [51]. When subjected to low frequency (0.01 Hz) and strain magnitude (1.4%), in vitro human breast cancer cells were shown to increase their length and form filopodia in the initial stretching cycle but to undergo apoptosis in later stretching cycles [52].

Many studies have reported that the interactions (mechanical and chemical) between epithelial WT and transformed cells drive apical extrusion in a non-cell-autonomous fashion [14,50,53]. The importance of the Ras^V12^-WT heterotypic interactions for the cellular behaviour of Ras^V12^ cells is further supported by our result that surrounding WT cells guide the Ras^V12^ protrusion growth direction. The apical extrusion process is known to involve the generation and transmission of forces via the cytoskeleton and cell-cell adhesions [14,26,27]. Overall, our study shows that strains affect key mechanical structures differently in WT than in Ras^V12^ cells, including the cortical actin, e-cadherin, basal SFs, cell body orientation, and vinculin. Strain-induced perturbations bring the cellular system to a new mechanical configuration. The achieved state, at least partially regulated by the Rho-ROCK pathway, favors basal invasion of the epithelia by the Ras^V12^ cells rather than apical extrusion. Better understanding of how these mechanically-induced reorganizations are orchestrated may help in finding new potential avenues for preventing cancer spreading, for instance by targeting mechanical strain pathways. Our results also show that pharmaceutical treatments can have an opposite effect on the Ras^V12^ cell’s behavior, depending if the system is being stretched or not. Of course, we must recognize that while complex biophysical phenomena were observed in this study, the system being examined is essentially a proof-of-concept model for transformed cell extrusion and invasion phenotypes. Certainly, much more work in vitro and in vivo will be required to understand how the phenomena reported here translate into more complex systems and animal study. However, these insights gained here demonstrate the importance of considering the implementation of mechanically dynamic environments when designing cancer research studies [54], especially in the very earliest stages of therapy development.

## 4 Materials and Methods

### 4.1 Microfluidic device and strain characterization

A detailed description of the microfluidic device fabrication and characterization can be found in previously reported work[29]. Briefly, the PDMS (Sylgard184, Ellsworth Adhesives Canada Corporation) microfluidic stretchers were prepared using standard microfabrication techniques, including UV photolithography and soft lithography. Each device contains a thin square suspended PDMS membrane (1.6 mm x 1.6 mm x 10 μm) that is attached to four vacuum chambers (two were used here). The cyclic action of a pair of isolated vacuum chambers (sinusoidal wave form, controlled via a homemade Labview program) deforms the walls in which the membrane is anchored, thus stretching the latter. The generated strain field pattern is biologically relevant with strain amplitudes varying from 3 to 9 % and strain gradient amplitudes varying from 0 to 12 % mm^-1^, mimicking in vivo strain non-uniformity[55]. The estimated stiffness of the PDMS membrane is 0.5 MPa based on previously reported characterizations of similar PDMS substrates [56,57].

### 4.2 Cell lines, cell culture, and inhibitors

Ras^V12^ is a stable MDCK cell line expressing in a tetracycline-inducible manner the oncogene Ras (GFP-tagged) [13]. Prior to seeding the cells into the microdevice, the PDMS membrane was air-plasma treated (Glow Research, Tempe, AZ, USA) at 70 W for 5 minutes. Sterilization and washing were performed by flowing 70% ethanol for 5 minutes followed by autoclaved phosphate buffered saline (PBS) solution for another 5 minutes. The membrane was then coated with Rat-tail collagen I (5 μg cm^-2^, Gibco) for 4 hours and rinses with PBS. A solution of suspended cells (ratio of 1:75 Ras^V12^ to WT MDCK) was injected into the device and incubated for 10 hours to grow the monolayer. Cyclic stretching of the well-adhered co-culture was initiated immediately after Ras^V12^ activation with tetracycline (2μg/ml). The monolayer was stretched for 24 hours at 1 Hz in the incubator. Inhibition studies of focal adhesion kinase (PF-573228, 10 µM, Selleckchem Inhibitor Expert, catalogue no. S2013), Rho-kinase (Y-27632; 10 µM, S cadherin ma, catalogue no. Y0503), and myosin-II (blebbistatin; 10 µM, Sigma, catalogue no. B0560), were achieved by exposing the cells immediately before stretching. The pharmacological treatments used were stored in dimethyl sulfoxide (DMSO) stock solutions. The drug-free experiments were supplemented with 0.1% DMSO.

### 4.3 Immunofluorescence Microscopy

Prior to DNA, actin, and e-cadherin staining, cells were fixed with 3.5% paraformaldehyde (15 min) and permeabilized with Triton X-100 (3 min) at 37°C. The DNA was labelled with DAPI (Invitrogen, catalogue no. D1306), the actin with phalloidin conjugated to Alexa Fluor 546 (Invitrogen, catalogue no. A22283), and the e-cadherin with monoclonal rat anti-e-cadherin (Sigma, catalogue no. U3254) primary antibody followed by polyclonal anti-rat IgG secondary antibody conjugated CF-647 (Sigma, catalogue no. SAB4600186). On-chip cell imaging was carried on with a long-distance 25x objective (NA=1.1) and an upright laser scanning multiphoton confocal microscope (Nikon **A1RsiMP**). The fluorescence signal of the images presented here have been optimized by adjusting the brightness/contrast settings using the program ImageJ. The despeckle function has only been used on images showing large cell components to remove salt and pepper noise. The maximum intensity z-projection of either the full stack or of a subset of slices is shown depending on the element of interest. Further details are provided in section S.2 of the SI.

### 4.4 Data analysis

Statistical significances were determined using unpaired two-tailed Student’s *t* tests, assuming unequal variances. The image analysis was performed using homemade Matlab programs making use of the image processing toolbox. Details of the image analysis strategies, as well as additional information on the material and experimental methods, are provided in section S.2 of the SI.

## 6 Author Contributions

S.C.L., M.G., and A.E.P. have designed the research. S.C.L. has developed the experimental setups, stretching devices, and experimental protocols. S.C.L. has performed the experiments and imagining. S.C-L. and H.J-R. have both contributed to develop the analytic tools and analyzed the data. Finally, all four authors have contributed to writing or editing the manuscript.

## 7 Funding and Acknowledgments

This work was supported by individual Natural Sciences and Engineering Research Council (NSERC) Discovery Grants to M.G. and A.E.P. S.C.-L. was supported by NSERC Postgraduate Scholarships-Doctoral (PGS D). M.G. acknowledges support from the Ontario Ministry of Research and Innovation, and the Canada Foundation for Innovation (CFI). A.E.P. gratefully acknowledges generous support from the Canada Research Chairs (CRC) program. We thank professor Yasuyuki Fujita (Hokkaido University) for providing the Ras^V12^ cell line.

## References

1. Gudipaty SA, Rosenblatt J. Epithelial cell extrusion: Pathways and pathologies. Semin Cell Dev Biol (2017) 67:132–140. doi:10.1016/j.semcdb.2016.05.010

2. Guillot C, Lecuit T. Mechanics of Epithelial Tissue Homeostasis and Morphogenesis. Science (80-) (2013) 340:1185–1189. doi:10.1126/science.1235249

3. Marchiando AM, Graham WV, Turner JR. Epithelial Barriers in Homeostasis and Disease. Annu Rev Pathol Mech Dis (2010) 5:119–144. doi:10.1146/annurev.pathol.4.110807.092135

4. Kuipers D, Mehonic A, Kajita M, Peter L, Fujita Y, Duke T, Charras G, Gale JE. Epithelial repair is a two-stage process driven first by dying cells and then by their neighbours. J Cell Sci (2014) 127:1229–1241. doi:10.1242/jcs.138289

5. Rosenblatt J, Raff MC, Cramer LP. An epithelial cell destined for apoptosis signals its neighbors to extrude it by an actin- and myosin-dependent mechanism. Curr Biol (2001) 11:1847–1857. doi:10.1016/S0960-9822(01)00587-5

6. Harris AR, Peter L, Bellis J, Baum B, Kabla AJ, Charras GT. Characterizing the mechanics of cultured cell monolayers. Proc Natl Acad Sci (2012) 109:16449–16454. doi:10.1073/pnas.1213301109

7. Slattum GM, Rosenblatt J. Tumour cell invasion: an emerging role for basal epithelial cell extrusion. Nat Rev Cancer (2014) 14:495–501. doi:10.1038/nrc3767.Tumour

8. Kajita M, Fujita Y. JB Review EDAC□: Epithelial defence against cancer — cell competition between normal and transformed epithelial cells in mammals. J Biochem (2015) 158:15–23. doi:10.1093/jb/mvv050

9. Bos JL. ras Oncogenes in Human Cancer□: A Review ras Oncogenes in Human Cancer□: A Review1. (1989) 4682–4689.

10. Rajalingam K, Schreck R, Rapp UR, Albert ??tefan. Ras oncogenes and their downstream targets. Biochim Biophys Acta - Mol Cell Res (2007) 1773:1177–1195. doi:10.1016/j.bbamcr.2007.01.012

11. Xia M, Land H. Tumor suppressor p53 restricts Ras stimulation of RhoA and cancer cell motility. Nat Struct Mol Biol (2007) 14:215–223. doi:10.1038/nsmb1208

12. Zondag GCM, Evers EE, Ten Klooster JP, Janssen L, Van Der Kammen RA, Collard JG. Oncogenic Ras downregulates Rac activity, which leads to increased Rho activity and epithelial-mesenchymal transition. J Cell Biol (2000) 149:775–781. doi:10.1083/jcb.149.4.775

13. Hogan C, Dupré-Crochet S, Norman M, Kajita M, Zimmermann C, Pelling AE, Piddini E, Baena-López LA, Vincent J-P, Itoh Y, et al. Characterization of the interface between normal and transformed epithelial cells. Nat Cell Biol (2009) 11:460–467. doi:10.1038/ncb1853

14. Kajita M, Sugimura K, Ohoka A, Burden J, Suganuma H, Ikegawa M, Shimada T, Kitamura T, Shindoh M, Ishikawa S, et al. Filamin acts as a key regulator in epithelial defence against transformed cells. Nat Commun (2014) 5:1–13. doi:10.1038/ncomms5428

15. Slattum G, Gu Y, Sabbadini R, Rosenblatt J. Autophagy in oncogenic K-Ras promotes basal extrusion of epithelial cells by degrading S1P. Curr Biol (2014) 24:19–28. doi:10.1016/j.cub.2013.11.029

16. Gu Y, Shea J, Slattum G, Firpo MA, Alexander M, Golubovskaya VM, Rosenblatt J. Defective apical extrusion signaling contributes to aggressive tumor hallmarks. Elife (2015) 2015:1–17. doi:10.7554/eLife.04069

17. Marshall TW, Lloyd IE, Delalande JM, Nathke I, Rosenblatt J. The tumor suppressor adenomatous polyposis coli controls the direction in which a cell extrudes from an epithelium. Mol Biol Cell (2011) 22:3962–3970. doi:10.1091/mbc.E11-05-0469

18. Bravo-Cordero JJ, Hodgson L, Condeelis JS. Spatial regulation of tumor cell protrusions by RhoC. Cell Adh Migr (2014) 8:263–267. doi:10.4161/cam.28405

19. Humphrey JD, Dufresne ER, Schwartz M a. Mechanotransduction and extracellular matrix homeostasis. Nat Rev Mol Cell Biol (2014) 15:802–812. doi:10.1038/nrm3896

20. Kim EH, Oh N, Jun M, Ko K, Park S. Effect of cyclic stretching on cell shape and division. BioChip J (2015) 9:306–312. doi:10.1007/s13206-015-9406-x

21. Jain RK, Martin JD, Stylianopoulos T. The role of mechanical forces in tumor growth and therapy. Annu Rev Biomed Eng (2014) 16:321–346. doi:10.1146/annurev-bioeng-071813-105259.The

22. Kumar S, Weaver VM. Mechanics, malignancy, and metastasis: The force journey of a tumor cell. Cancer Metastasis Rev (2009) 28:113–127. doi:10.1007/s10555-008-9173-4.Mechanics

23. Pickup MW, Mouw JK, Weaver VM. The extracellular matrix modulates the hallmarks of cancer. EMBO Rep (2014) 15:1243–1253. doi:10.15252/embr.201439246

24. Menon S, Beningo KA. Cancer cell invasion is enhanced by applied mechanical stimulation. PLoS One (2011) 6: doi:10.1371/journal.pone.0017277

25. Gasparski AN, Ozarkar S, Beningo KA. Transient mechanical strain promotes the maturation of invadopodia and enhances cancer cell invasion in vitro. J Cell Sci (2017) 130:1965–1978. doi:10.1242/jcs.199760

26. Wu SK, Lagendijk AK, Hogan BM, Gomez GA, Yalp AS. Active contractility at E-cadherin junctions and its implications for cell extrusion in cancer. Cell Cycle (2015) 14:315–322.

27. Michael M, Lefevre JG, Parton RG, Wu SK, Gomez GA, Michael M, Verma S, Cox HL, Lefevre JG, Parton RG, et al. Cortical F-actin stabilization generates apical-lateral patterns of junctional contractility that integrate cells into …; Cortical F-actin stabilization generates apical – lateral patterns of junctional contractility that integrate cells into epithelia. Nat Cell Biol (2014) 16:1–15. doi:10.1038/ncb2900

28. Lu D, Kassab GS. Role of shear stress and stretch in vascular mechanobiology. J R Soc Interface (2011) 8:1379–1385. doi:10.1098/rsif.2011.0177

29. Chagnon-Lessard S, Jean-Ruel H, Godin M, Pelling AE. Cellular orientation is guided by strain gradients. Integr Biol (2017) 9:607–618. doi:10.1039/C7IB00019G

30. Goldmann WH, Fabry B, Klemm AH, Kienle S, Scha TE. Focal Adhesion Kinase Stabilizes the Cytoskeleton. (2011) 101:2131–2138. doi:10.1016/j.bpj.2011.09.043

31. Wang HB, Dembo M, Hanks SK, Wang Y. Focal adhesion kinase is involved in mechanosensing during fibroblast migration. Proc Natl Acad Sci U S A (2001) 98:11295–300. doi:10.1073/pnas.201201198

32. Zhou J, Aponte-Santamaría C, Sturm S, Bullerjahn JT, Bronowska A, Gräter F. Mechanism of Focal Adhesion Kinase Mechanosensing. PLoS Comput Biol (2015) 11:1–16. doi:10.1371/journal.pcbi.1004593

33. Tomakidi P, Schulz S, Proksch S, Weber W, Steinberg T. Focal adhesion kinase (FAK) perspectives in mechanobiology: implications for cell behaviour. Cell Tissue Res (2014) 357:515–526. doi:10.1007/s00441-014-1945-2

34. McLean GW, Carragher NO, Avizienyte E, Evans J, Brunton VG, Frame MC. The role of focal-adhesion kinase in cancer — a new therapeutic opportunity. Nat Rev Cancer (2005) 5:505–515. doi:10.1038/nrc1647

35. Winklbauer R. Cell adhesion strength from cortical tension – an integration of concepts. (2015) 2:3687–3693. doi:10.1242/jcs.174623

36. Wodarz a, Nathke I. Cell polarity in development and cancer. Nat Cell Biol (2007) 9:1016–1024. doi:ncb433 [pii]\n10.1038/ncb433

37. Cowin P, Rowlands TM, Hatsell SJ. Cadherins and catenins in breast cancer. Curr Opin Cell Biol (2005) 17:499–508. doi:10.1016/j.ceb.2005.08.014

38. Cavallaro U, Christofori G. Cell adhesion and signalling by cadherins and Ig-CAMs in cancer. Nat Rev Cancer (2004) 4:118–132. doi:10.1038/nrc1276

39. Chang Y-WE, Jakobi RRB and R. Targeting RhoA/Rho Kinase and p21-Activated Kinase Signaling to Prevent Cancer Development and Progression. Recent Pat Anticancer Drug Discov (2009) 4:110–124. doi:http://dx.doi.org/10.2174/157489209788452830

40. Marjoram RJ, Lessey EC, Burridge K. Regulation of RhoA Activity by Adhesion Molecules and Mechanotransduction. Curr Mol Med (2014) 14:199–208. doi:10.2174/1566524014666140128104541

41. Sahai E, Marshall CJ. ROCK and Dia have opposing effects on adherens junctions downstream of Rho. Nat Cell Biol (2003) 4: doi:10.1038/ncb796

42. Lecuit T, Lenne P-F. Cell surface mechanics and the control of cell shape, tissue patterns and morphogenesis. Nat Rev Mol Cell Biol (2007) 8:633–644.

43. Ueda S, Blee AM, Macway KG, Renner DJ, Yamada S. Force dependent biotinylation of myosin IIA by α-catenin tagged with a promiscuous biotin ligase. PLoS One (2015) 10:1–15. doi:10.1371/journal.pone.0122886

44. Abiko H, Fujiwara S, Ohashi K, Hiatari R, Mashiko T, Sakamoto N, Sato M, Mizuno K. Rho guanine nucleotide exchange factors involved in cyclic-stretch-induced reorientation of vascular endothelial cells. J Cell Sci (2015) 128:1683–95. doi:10.1242/jcs.157503

45. Bays JL, DeMali KA. Vinculin in cell–cell and cell–matrix adhesions. Cell Mol Life Sci (2017) 74:2999–3009. doi:10.1007/s00018-017-2511-3

46. Yamaguchi H, Condeelis J. Regulation of the actin cytoskeleton in cancer cell migration and invasion. Biochim Biophys Acta - Mol Cell Res (2007) 1773:642–652. doi:10.1016/j.bbamcr.2006.07.001

47. Bravo-Cordero JJ, Hodgson L, Condeelis J. Directed cell invasion and migration during metastasis. Curr Opin Cell Biol (2012) 24:277–283. doi:10.1016/j.ceb.2011.12.004

48. Alibert C, Goud B, Manneville JB. Are cancer cells really softer than normal cells? Biol Cell (2017) 109:167–189. doi:10.1111/boc.201600078

49. Struckhoff AP, Rana MK, Worthylake RA. RhoA can lead the way in tumor cell invasion and metastasis. Front Biosci (2011) 16:1915–26.

50. Yamamoto S, Yako Y, Fujioka Y, Kajita M, Kameyama T, Kon S, Ishikawa S, Ohba Y, Ohno Y, Kihara A, et al. A role of the sphingosine-1-phosphate (S1P)-S1P receptor 2 pathway in epithelial defense against cancer (EDAC). Mol Biol Cell (2016) 27:491–499. doi:10.1091/mbc.E15-03-0161

51. Berrueta L, Bergholz J, Munoz D, Muskaj I, Badger GJ, Shukla A, Kim HJ, Zhao JJ, Langevin HM. Stretching Reduces Tumor Growth in a Mouse Breast Cancer Model. Sci Rep (2018) 8:1–7. doi:10.1038/s41598-018-26198-7

52. Yadav S, Vadivelu R, Ahmed M, Barton M, Nguyen NT. Stretching cells – An approach for early cancer diagnosis. Exp Cell Res (2019) 378:191–197. doi:10.1016/j.yexcr.2019.01.029

53. Hogan C, Dupré-crochet S, Norman M, Kajita M, Zimmermann C, Pelling E, Piddini E, Baena-lópez LA, Vincent J, Itoh Y, et al. Characterization of the interface between normal and transformed epithelial cells. Nat Cell Biol (2009) 11:460–7. doi:10.1038/ncb1853

54. Mei X, Middleton K, Shim D, Wan Q, Xu L, Ma YV, Devadas D, Walji N, Wang L, Young EWK, et al. Microfluidic platform for studying osteocyte mechanoregulation of breast cancer bone metastasis. Integr Biol (2019) 11:119–129. doi:10.1093/intbio/zyz008

55. Richardson WJ, Metz RP, Moreno MR, Wilson E, Moore JE. A Device to Study the Effects of Stretch Gradients on Cell Behavior. J Biomech Eng (2011) 133:101008(1–9). doi:10.1115/1.4005251

56. Eddington DT, Crone WC, Beebe DJ. Development of process protocols to fine tune polydimethylsiloxane material properties. 7th lnternational Conf Miniaturized Chem Blochemlcal Anal Syst (2003) 7:1089–92. doi:10.1007/s10544-005-3029-2

57. Park JY, Yoo SJ, Lee EJ, Lee DH, Kim JY, Lee SH. Increased poly(dimethylsiloxane) stiffness improves viability and morphology of mouse fibroblast cells. Biochip J (2010) 4:230–236. doi:10.1007/s13206-010-4311-9

