## Supplementary Material for "Mechanotransduction of strain regulates an invasive phenotype in newly transformed epithelial cells"

### S.1 ADDITIONAL FIGURES AND MOVIE

**Movie M1:** A PDMS pneumatic-based microfluidic stretcher is cyclically stretched. The action of the vacuum pumps pulls a thin PDMS membrane on which a MCDK monolayer is cultured. The size of the cell chamber is 1.6 mm x 1.6 mm.

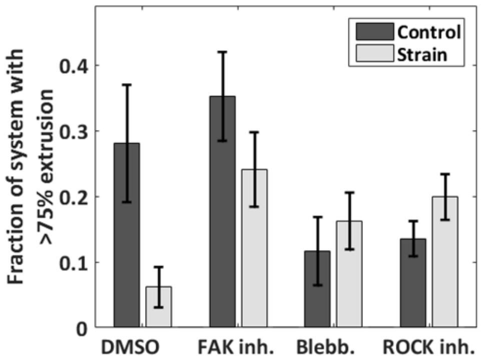


**Fig. S1.** Effect of mechanical strain on $Ras^{V12}$ apical extrusion. The apical extrusion is quantified here by the fraction of the $Ras^{V12}$ cell clusters in which at least 75% of the cells are above the average surrounding WT height. In addition to the drug-free case (DMSO), three drugs were tested: FAK inhibitor, blebbistatin, and ROCK inhibitor. From left to right: n = 234, 188, 78, 139, 268, 233, 237 and 334 cells each from 3 or 4 independent experiments. Data are mean ± s.e.m.

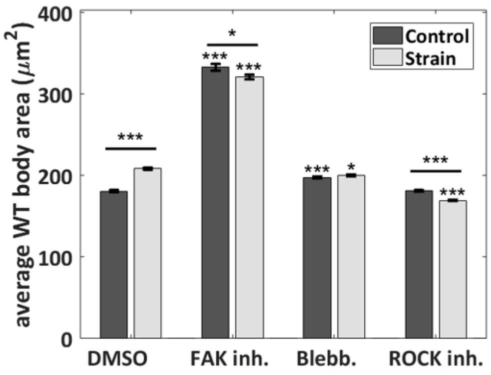


**Fig. S2.** Effect of mechanical strain on the WT cell body size ($\mu m^2$). In addition to the drug-free case (DMSO), three drugs were tested: FAK inhibitor, blebbistatin, and ROCK inhibitor. Data are mean ± s.e.m. *** $P<0.0001$, **$P<0.005$, *$P<0.05$. Stars immediately above the individual bars relate to the corresponding DMSO control or DMSO strain condition. Stars above the horizontal lines refer to the significance between the control and strain data. For each bar, n>2000 from 3 to 4 independent experiments.

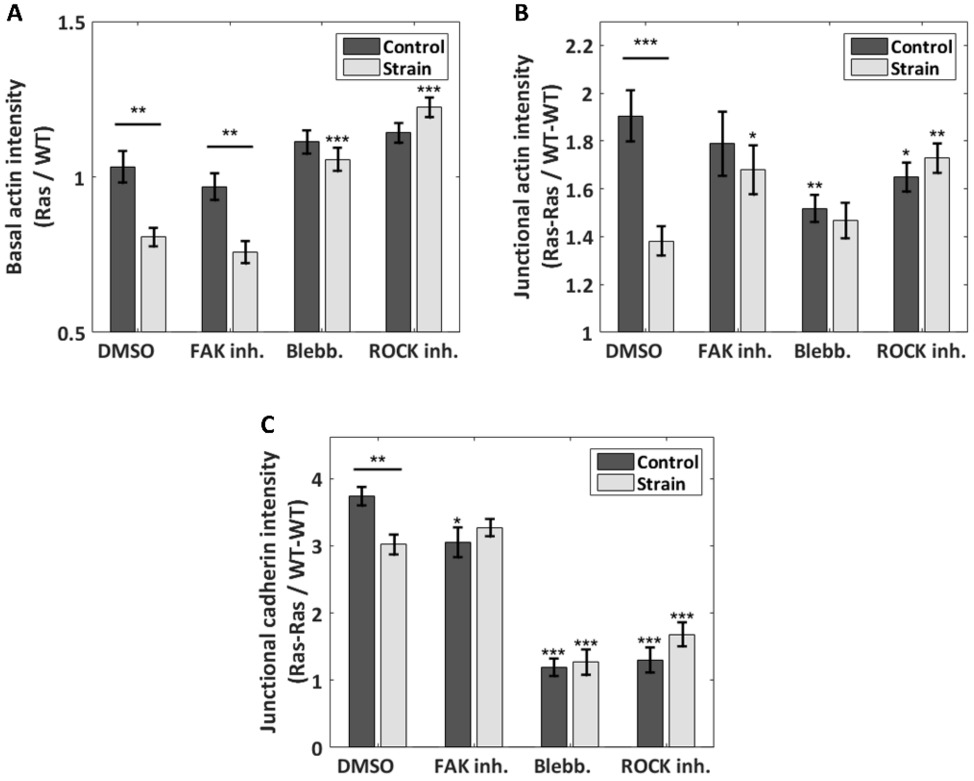


**Fig. S3.** Effect of mechanical strain on different actin and e-cadherin ratios. (A) Ratio of the basal actin intensity, $Ras^{V12}$/ WT cells. (B) Ratio of the junctional actin intensity, $Ras^{V12}$- $Ras^{V12}$ interface over WT-WT interface. (C) Ratio of the junctional e-cadherin intensity, $Ras^{V12}$- $Ras^{V12}$ interface over WT-WT interface. In addition to the drug-free case (DMSO), three drugs were tested: FAK inhibitor, blebbistatin, and ROCK inhibitor. Data are mean ± s.e.m. *** $P<0.0001$, **$P<0.005$, *$P<0.05$. Stars immediately above the individual bars relate to the corresponding DMSO control or DMSO strain condition. Stars above the horizontal lines refer to the significance between the control and strain data. From left to right, n = 43, 48, 29, 44, 56, 56, 53, and 57 images, each from 3 to 4 independent experiments. The average ratios of each image were compiled and they were used to determine the mean and the s.e.m. of the data points reported; the total number of $Ras^{V12}$ cells contained in each set of images is given in the caption of Fig. S1 (above).

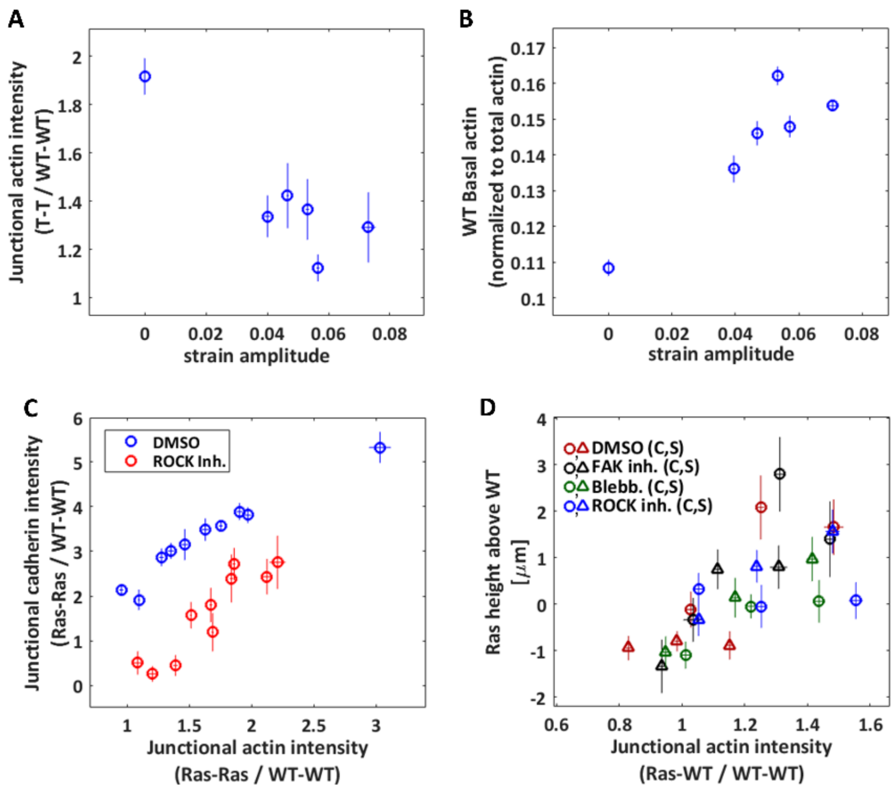


**Fig. S4.** Junctional intensity of different cell components. (A) Junctional actin intensity, $Ras^{V12}$–$Ras^{V12}$ interface over WT - WT interface, as a function of the local strain amplitude, for the DMSO (drug-free) condition. (B) WT basal actin intensity as a function of the local strain amplitude, for the DMSO (drug-free) condition. For(A) and (B), the zero-strain points correspond to the average of the control data, and the non-zero strain points were obtained by binning the data of the stretched experiments into five bins each containing an equal (37-38) number of cells. (C) Junctional e-cadherin intensity, $Ras^{V12}$- $Ras^{V12}$ interface over WT-WT interface, as a function of the junctional actin intensity, $Ras^{V12}$- $Ras^{V12}$ interface over WT-WT interface, for the DMSO (drug-free) and ROCK inhibitor conditions. The points were obtained by binning the data from the control and stretching experiments each into 5 bins containing an equal number of cells (37-67). (D) Correlation between $Ras^{V12}$ extrusion height and the junctional actin intensity ratio of $Ras^{V12}$- WT interface over WT-WT interface. The data from each of the 8 conditions (DMSO and three drugs, static (c) and strain (s)) were separated in three bins each of equal number of cells. $n_{control\_DMSO} = 234$, $n_{strain\_DMSO} = 188$, $n_{control\_FAK} = 78$, $n_{strain\_FAK} = 139$, $n_{control\_blebb} = 268$, $n_{strain\_blebb} = 233$, $n_{control\_ROCK} = 237$, and $n_{strain\_ROCK} = 334$ cells each from 3 or 4 independent experiments. Data are mean ± s.e.m.

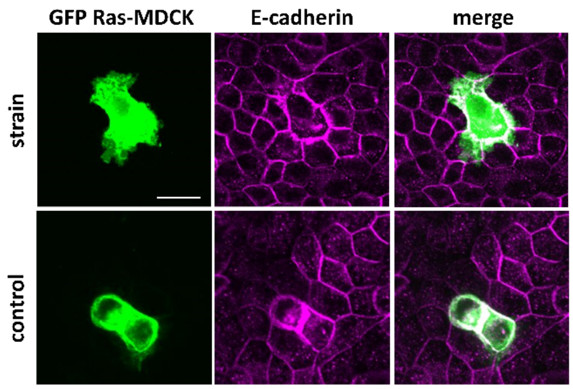


**Fig. S5.** Representative confocal images of the $Ras^{V12}$-WT culture showing a GFP-$Ras^{V12}$ cells (green) and e-cadherin (magenta) for the control and the strain DMSO conditions. Scale bar is 25 µm.

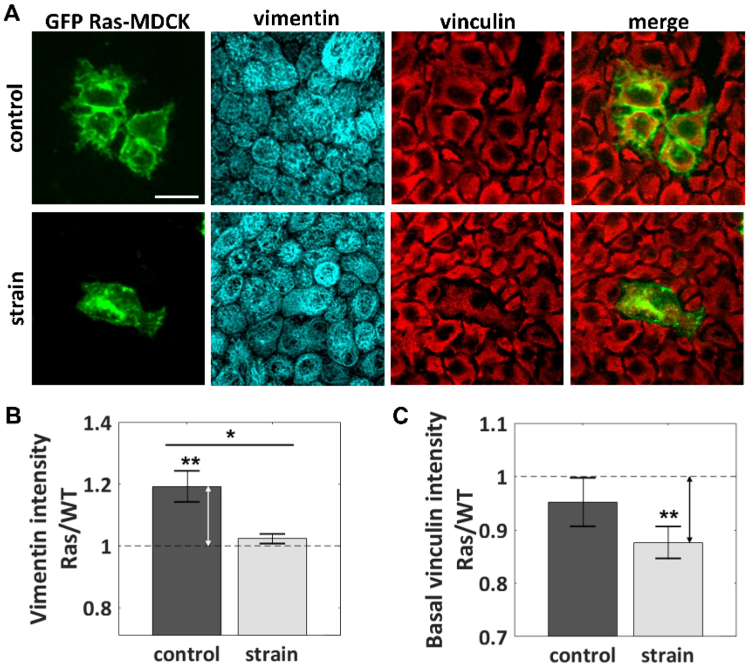


**Fig. S6.** Representative confocal fluorescence images of the $Ras^{V12}$-WT MDCK culture showing GFP-$Ras^{V12}$ cells (green), vimentin (cyan), and basal vinculin (red) for the control and the strain DMSO conditions (A). Scale bar is 30 µm. (B) Ratio of the vimentin intensity, $Ras^{V12}$ cells over WT cells, for the control and the strain DMSO conditions. (C) Ratio of the vinculin basal intensity, $Ras^{V12}$ cells over WT cells, for the control and the strain DMSO conditions. (B, C) Data are mean ± s.e.m. *$P<0.05$, **$P<0.005$. Stars immediately above the individual bars refer to the statistical significance between the ratio observed and a ratio of 1. Stars above the horizontal lines refer to the significance between the control and strain data. The number of control and strain images that were analyzed were $n_{control}=16$ and $n_{strain}=18$, each from 3 independent experiments.

### S.2 Materials and Methods Additional Information

**Staining Procedures**:

| Cell structures | Excitation λ [nm] | Fixation method | Staining method | Product ID |
|---|---|---|---|---|
| DNA | 358 | PFA or Methanol | DAPI | Invitrogen, catalogue no. D1306 |
| Actin | 546 | PFA | Phalloidin conjugated to Alexa Fluor 546 (1:100 dilution for 30 min) | Invitrogen, catalogue no. A22283 |
| E-cadherin | 647 | PFA | **1)** Monoclonal anti-e-cadherin primary antibody produced in rat (1:1000 dilution for 30 min)<br>**2)** Polyclonal anti-rat IgG (H+L) secondary antibody conjugated CF-647 produced in goat (1:200 dilution for 30 min) | **1)** Sigma, catalogue no. U3254<br>**2)** Sigma, catalogue no. SAB4600186 |
| Vinculin | 546 | Methanol | **1)** Monoclonal anti-vinculin primary antibody produced in mouse (1:200 dilution for 30 min)<br>**2)** Anti-mouse IgG secondary antibody conjugated to Alexa Fluor 546 fluorophore produced in goat (1:200 dilution for 30 min) | **1)** Sigma, catalogue no. V9131<br>**2)** Invitrogen, catalogue no. A11003 |
| Vimentin | 647 | Methanol | **1)** Monoclonal anti-vimentin primary antibody produced in rabbit (1:500 dilution for 30 min)<br>**2)** Anti-Rabbit IgG (H+L) secondary antibody conjugated to Alexa Fluor Plus 647 (1:200 dilution for 30 min) | **1)** Abcam, catalogue no. AB92547<br>**2)** Invitrogen, catalogue no. A32733 |
| GFP-Ras | 488 | PFA or Methanol | NA | NA |

**Table S1. Staining procedure summary.** The PFA fixation method consists in employing 3.5% paraformaldehyde for 15 min at 37°C and Triton X-100 for 3 min for permeabilization. The methanol method consists in employing 95% methanol at -20°C for 3 min. Both of these methods are followed by 3 washing steps and 15 min incubation with PBS. Furthermore, following each staining step, the cells were washed 3 times and incubated for 15 min with blocking buffer [5% horse serum (Sigma) in phosphate buffered saline (PBS)] for actin, e-cadherin, vinculin, and with PBS only for DNA and vimentin. The antibodies were diluted in blocking buffer. Following the staining procedure, the devices were filled with PBS to maintain cell hydration before imaging. Standard staining protocols were adapted to suit the microdevice environment.

**Membrane strain characterization:**

The strain field was locally determined by employing a point-by-point characterization method. Fluorescence beads (FluoSpheres, 200 nm, Invitrogen) were incorporated in the PDMS membrane and imaged for different stretched states (10 states) with an EPI fluorescence microscope (Nikon, Tokyo, Japan). A homemade Matlab program was generated to track the bead displacements and calculate the Green strain matrix elements.

**Cell culture:**

MDCK and MDCK-$Ras^{V12}$ transformed epithelial cells were cultured in Dulbecco's Modified Eagle's Medium (DMEM) supplemented with 10% FBS, 50mg/ml streptomycin, and 50U/ml penicillin antibiotics (Hyclone Laboratories, Logan, UT, USA). All the cells were cultured in a standard incubator at 37°C with 5% $CO_2$, initially on d=100 mm tissue culture dishes (Corning) and then on the microfluidic stretcher membranes.

**Data processing and analysis:**

For each $Ras^{V12}$ cell, 3-dimensional fluorescence imaging was performed with a Z-step size of 1 μm. Each stack typically contained between 15 and 40 slices, depending the $Ras^{V12}$ cell extrusion level. Automated image analysis programs were developed in Matlab, taking advantage of the image processing toolbox. First, in each image, the X-Y-Z nuclei positions were extracted, the cell contours were determined, and the $Ras^{V12}$ cell bodies were identified. Visual inspection of each of these steps was carried out for every image, and manual corrections were performed when necessary. The majority of the subsequent analysis built upon this extraction. Specifically, the local stacks considered for the basal (-5 to -3 μm), mid-cell (-1 to +1 um), and upper-cell (>1 um) Z-slice projections were determined based on the local nuclei average height ("0-height"). The protrusion areas were determined based on the basal level GFP signal outside of the $Ras^{V12}$ cell bodies. The height difference between the $Ras^{V12}$ cells and the surrounding WT cells was determined from the apical maxima of the Z-profiles of the actin intensity averaged over the cell areas. The junctional regions were identified based on the extracted cell contours, considering 11 pixel (~4 μm) wide regions centered on the cell frontiers. The actin and e-cadherin channels were summed over all Z-slices for determining the average junctional intensity ratios in Fig. 2B,G, Fig. S3B,C, and Fig. S4A,C,D. The actin channel was summed over the local basal slices for determining the basal actin in Fig. 2D,E, Fig. S3A, and Fig. S4B . For Fig. 3B, the orientation of the protrusion segments and basal actin SFs were both determined manually on different identification sessions. This was achieved by displaying in each case only the element to be characterized in that session (to prevent any bias). To compile the histogram in Fig. 4D, only the WT SFs which were outside a protrusion and which approached its major axis within 4 μm or less were considered. The actin channel was summed over the local upper-cell Z-slices in Fig. 3C (to exclude basal SFs). The regionprops function of Matlab was used to determine the cell body area in Fig. S2, based on the cell contour extraction. Finally, for the vinculin and vimentin analysis of Fig. S6, the $Ras^{V12}$ cell regions were determined based on the

GFP channel alone (thus including both the protrusions and the cell bodies). The vinculin channel was summed over the local basal slices while the vimentin was summed over all Z-slices. Additional details are presented in the table S2 (below).

**Table S2. Image analysis summary.**

<table>
<tr><th>Description</th><th>Example of output image<br>(DMSO control, static state)</th></tr>
<tr><td>Convert confocal stacks into a 4-dimensional Matlab array.</td><td>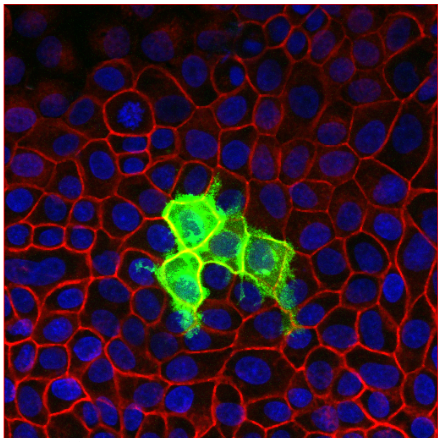<br>GFP-Ras$^{V12}$ cells in green, junctional actin in red, and nuclei in blue</td></tr>
<tr><td>Extract nuclei positions (X,Y,Z):<br>• Isolate the DAPI channel, extract binary images for each Z-slice, and determine X,Y center of the identified particles;<br>• Go up slice-by-slice to identify and correct nuclei which were erroneously merged together into a single particle; repeat in the other direction;<br>• Regroup nuclei identified from different Z-slices which belong to the same particle;<br>• Determine the average X and Y positions, and find the Z position based on the Z intensity profile of a small region around the corresponding X,Y center;<br>• Inspect image and “manually” remove or add nuclei if necessary;</td><td>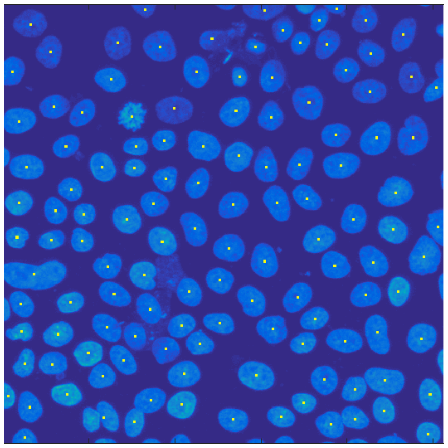<br>Identified nuclei shown with yellow dots</td></tr>
</table>

<table>
<tr>
<td>

**Extract cell contours:**

- Isolate the actin channel and extract the projection of three stacks centered on the local average nuclei height to obtain optimal cell divisions based on the junctional actin signal;
- Extract binary image and perform watershed to obtain the first iteration of the cell contours;
- Scan through each newly identified cell;
  - if a cell contains no nucleus, reattribute its area to neighboring cells based on the actin profile and redefine cell contours;
  - if a cell contains more than one nucleus, split it based on the actin profile and redefine cell contours;
- Inspect image and "manually" correct cell divisions if necessary;

</td>
<td>

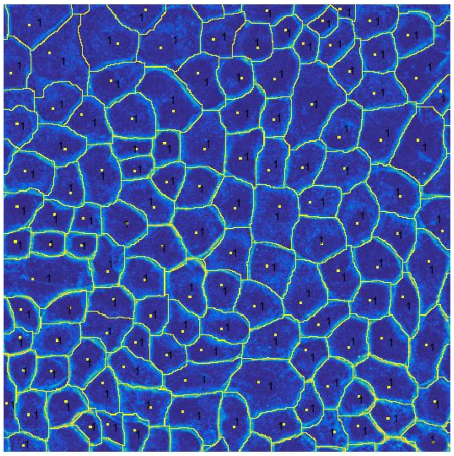

Extracted cell contours shown in yellow (overlaid on the junctional actin signal in cyan), identified nuclei shown with yellow dots, 1 means there is 1 nucleus in the identified cell.

</td>
</tr>
<tr>
<td>

**Determine which cell bodies are Ras$^{V12}$ transformed cells:**

- Isolate the GFP channel and extract the projection of three stacks above the local average nuclei height to obtain the non-basal GFP profile (i.e. excluding the roots);
- Extract binary image and determine which cells (identified in the previous step) are significantly covered by the GFP signal; these are the transformed cells;
- Inspect image and "manually" correct transformed cells identification if necessary;

</td>
<td>

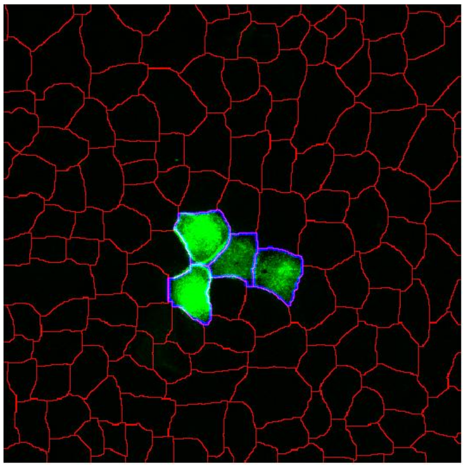

Identified cell contours in red, identified Ras$^{V12}$ cell contours in blue

</td>
</tr>
</table>

**Protrusion extraction:**

- Isolate the GFP channel and extract the projection of the stacks 3-5 μm below the local average nuclei height to obtain the basal GFP profile, which was found to provide the optimal view of the protrusions;
- Extract binary image of the basal GFP profile and compare it against the previously identified $Ras^{V12}$ cell contours: the GFP areas falling outside the transformed $Ras^{V12}$ cell body contours are considered to be protrusions;
- Associate each protrusion section to the proper transformed cell;

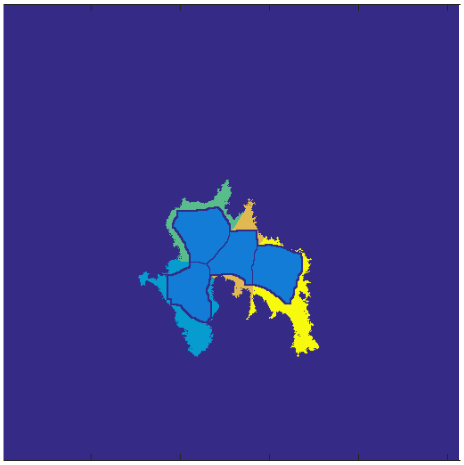

$Ras^{V12}$ cell bodies are in light blue and their protrusions are shown in other colors.

**Extraction of other parameters:**

- Following the identification of the nuclei heights, the cell contours, the $Ras^{V12}$ transformed cell bodies, and the $Ras^{V12}$ protrusions, all the other parameters can be extracted, such as:
  - Height difference between a $Ras^{V12}$ cell and the average surrounding WT cells, based on the apical maxima of the Z-profiles of the actin intensity (averaged in X-Y over the cell areas);
  - Area of the protrusions, normalized to the area of the corresponding $Ras^{V12}$ cell bodies;
  - WT-WT, $Ras^{V12}$-WT, and $Ras^{V12}$-$Ras^{V12}$ junctional actin or e-cadherin intensities, using the full Z-projection (see example of junctional region identification on the right);
  - WT and $Ras^{V12}$ basal actin and vinculin intensities, using the projection of the stacks 3 to 5 μm below the local average nuclei heights;
  - WT cell body orientations using Matlab's image processing toolbox to determine the major axis orientation;

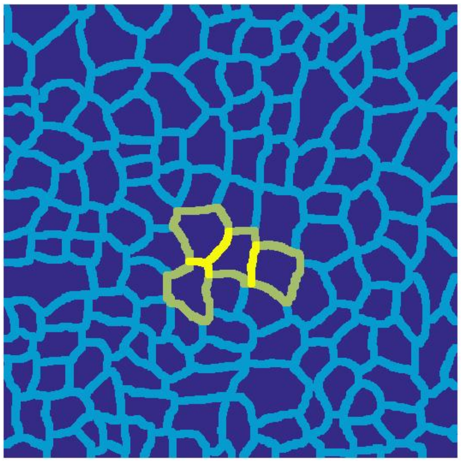

Light blue, green, and yellow regions are WT-WT, WT-$Ras^{V12}$, and $Ras^{V12}$-$Ras^{V12}$ junctions, respectively.
